# TKSM: Highly modular, user-customizable, and scalable transcriptomic sequencing long-read simulator

**DOI:** 10.1101/2023.06.12.544410

**Authors:** Fatih Karaoglanoglu, Baraa Orabi, Ryan Flannigan, Cedric Chauve, Faraz Hach

**Affiliations:** Computing Science Department, Simon Fraser University, 8888 University Dr, V5A 1S6, BC, Canada; Department of Computer Science,the University of British Columbia, 2366 Main Mall, V6T 1Z4, BC, Canada; Department of Urologic Sciences,the University of British Columbia, 2660 Oak Street, V6H 3Z6, BC, Canada; Vancouver Prostate Centre, 2660 Oak Street, V6H 3Z6, BC, Canada; Department of Mathematics, Simon Fraser University, 8888 University Dr, V5A 1S6, BC, Canada

**Author notes:** These authors contributed equally (Fatih Karaoglanoglu, Baraa Orabi).

## Abstract

**Motivation:** Transcriptomic long-read (LR) sequencing is an increasingly cost-effective technology for probing various RNA features. Numerous tools have been developed to tackle various transcriptomic sequencing tasks (e.g. isoform and gene fusion detection). However, the lack of abundant gold standard datasets hinders the benchmarking of such tools. Therefore, simulation of LR sequencing is an important and practical alternative to enable the assessment of these tools. While the existing LR simulators aim to imitate the sequencing machine noise and to target specific library protocols, they lack some important library preparation steps (e.g. PCR) and are difficult to modify to new and changing library preparation techniques (e.g. single-cell LRs).

**Results:** We present TKSM, a modular and scalable LR simulator. TKSM is designed so that each RNA modification step is targeted explicitly by a software module. This allows the user to assemble a simulation pipeline of any combination of TKSM modules to emulate the sequencing design the user is targeting. Additionally, the input/output of all the core modules of TKSM follow the same simple format (Molecule Description Format) allowing the user to easily extend TKSM with new modules targeting new library preparation steps.

**Availability:** TKSM is available as an open source software at https://github.com/vpc-ccg/tksm.

## 1 Introduction

Long-read (LR) sequencing technologies have become a cost-effective alternative to short-read (SR) sequencing for many genomic and transcriptomic sequencing tasks [Amarasinghe et al., 2020a]. LRs are shown to be useful for many transcriptomic tasks such as alternative isoform detection [Kovaka et al., 2019, Tang et al., 2020, Orabi et al., 2023], gene fusion detection [Liu et al., 2020, Karaoglanoglu et al., 2022], transcript-level expression analysis [Hu et al., 2021], or single-cell transcriptomic analysis [Tian et al., 2021, Ebrahimi et al., 2022, You et al., 2023].

However, due to the nature of LR sequencing as an emerging technology, there are very few well established benchmark datasets or gold standard datasets to assess transcriptomic LR bioinformatics tools. Such bioinformatics tools targeting these tasks require realistic simulations in order to assess their accuracy and performance. This includes the the simulation’s ability to explicitly target specific library or cellular processes such as single-cell barcoding and UMI tagging, PCR, or molecule truncation.

Existing LR simulators typically focus on simulating the sequencing process, i.e. the point of contact of sequencing platform with the RNA/DNA molecule [Wick, 2019, Li et al., 2020, Ono et al., 2022]. Some have extensions focusing on specific sequencing libraries such as transcriptomic, plasmid, or metagenomic samples [Hafezqorani et al., 2020, Yang et al., 2023]. However, existing tools are not designed with modularity in mind; they cannot be easily modified to address changes in the library preparation protocols such as adding a barcode tag or simulating the PCR process.

We describe TKSM, a software that simulates realistic transcriptomic long-read datasets. TKSM modular design allows to target a wide range of library/cell processes. The power of TKSM lays in two key aspects: i) the ease with which its simulation pipeline can be modified to cater to the user’s sequencing designs, ii) and its high performance in terms of time and memory use. TKSM is open source, accessible via GitHub.

## 2 Methods

TKSM is both flexible, in order that it can simulate a wide variety of datasets, and is extendable. It is composed of several independent modules, each representing a cellular (e.g. polyadenylation) or a library preparation (e.g. PCR) process that modifies a nucleic acid molecule. This design allows the user to simulate different sequencing protocols by using TKSM’s modules in various arrangements, imitating the different steps in the desired sequencing protocol. Additionally, this modular design allows TKSM to be easily extendable with future modules targeting library and cellular processes that we currently do not have modules for. To enable this modularity, we designed TKSM’s modules to take and generate files in the same format, that we call Molecule Description Format (MDF). An MDF file is a human-readable file that describes molecules by listing for each molecule its genomic intervals alongside any sequence-level modifications to these intervals (e.g. substitutions). The rationale for using a human-readable format is that the user can manually modify the intermediate files for their needs or write their own scripts that can generate or modify these MDF files. We expand on the details of MDF files in Section S1.4 of the Supplementary Materials. The only exceptions to this design pattern are the entry module which generates the initial set of molecules from a transcript abundance profile and the exit module which generates the reads obtained by simulating the sequencing of the given molecules.

Each of TKSM’s modules can be run as a separate process (`tksm <module_name>`). We also provide as part of TKSM a Snakemake [Mölder et al., 2021] script which can be configured by the user to specify a wide range of simulation experiments and run them all as a single command. Additionally, to optimize the computation time, we take advantage of Snakemake’s piped input/output feature to allow modules to start running the moment they receive any input from a previous module, rather than having to wait for the preceding module to terminate.

TKSM can use real sequencing datasets to parameterize the behaviour of its modules, or alternatively, these parameters can be specified manually by the user. For example, TKSM contains pre-processing modules to compute the expression profile of transcripts from a given real sample which is then used to generate the molecules in the initial MDF file, whose sequencing according to a chosen protocol will be simulated by the next modules.

## 2.1 TKSM modules

TKSM contains three classes of modules, defined by features of their input and output: i) entry-point modules start a TKSM pipeline and output an MDF file, ii) core modules take an MDF file as input and output another MDF file, and iii) exit (sequencing) modules take an MDF file as input and generates FASTA/FASTQ file(s) as output. Additionally, there are some preprocessing utilities in TKSM that can take a real sequencing dataset and output a model parameters for some of TKSM modules. A list of the implemented TKSM modules is presented in Figure 1. Additional modules and utilities can be implemented and easily integrated into TKSM in order to target steps in other sequencing protocols.

**Figure 1.**
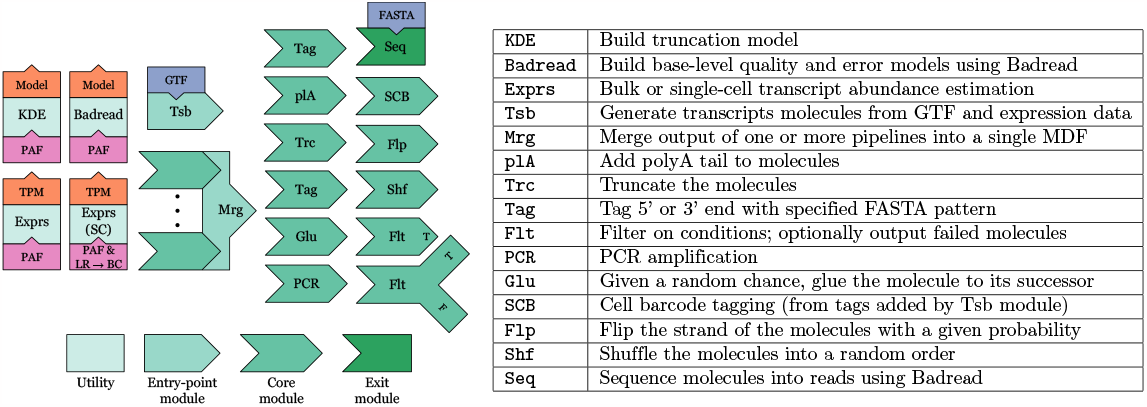
Existing TKSM modules and utilities alongside their high-level descriptions. TKSM is designed with modularity in mind; the user can specify simulation pipeline of their choosing by chaining any number of TKSM modules including the possibility using the same module multiple times.

### Abundance estimation utility

The abundance estimation utility takes as input a PAF file containing the alignment information of the long-reads of a real transcriptomic dataset to the transcriptome. We use Minimap2 [Li, 2018] to perform this transcriptomic mapping. Similarly to Trans-Nanosim [Hafezqorani et al., 2020], it computes the estimated abundance of different transcripts using an Expectation-Maximization (EM) algorithm implementation by Simpson [Simpson, 2018] which estimates transcript abundance from multi-mapping long-reads. This utility generates the transcript counts normalized by the total throughput and divided by one million (transcript per million or TPM).

#### Single cell expression

We augmented this abundance estimation method in order that it can also accept cellular barcode tags, generated by tools such as scTagger [Ebrahimi et al., 2022] or FLAMES [Tian et al., 2021]. This allows the generation of single-cell resolution transcript expression profiles, and thus to generate simulated single-cell datasets. To the best of our knowledge, TKSM is the only simulator that can generate single-cell transcriptomics long-read datasets. To do that, we maintain a separate count for each pair of cellular barcode and transcript. We then feed these counts to the standard EM algorithm for abundance estimation.

### Transcribing

The Transcribing module takes as input a gene annotation file (GTF) describing the genomic intervals corresponding to the transcripts and the transcript abundance estimation profile generated by TKSM Abundance estimation utility, and generates a set of molecules from which sequencing reads will be simulated. The Transcribing module is an entry-point module and thus it does not take an MDF file as input and it generates an MDF file as output. Given a user defined parameter *N*, the module samples *N* transcripts from the GTF file according to their abundance frequencies and outputs them in MDF format. Any cellular barcode information present in the abundance file for a given transcript is recorded in the output MDF.

#### Gene fusions

The Transcribing module can optionally generate gene fusion transcripts, for gene fusions induced by genomic structural variants. The user can specify genomic breakpoints of fusion-inducing structural variations (e.g. deletion, inversion or translocation); otherwise they are randomly chosen. Then, gene fusion transcripts are generated randomly from the transcripts of the involved genes and their expression (see details in Supplementary Material Section S1.6). As the MDF format enables the representation of various combinations of genomic intervals, irrespective of their chromosome or strand, and as downstream modules are indifferent to the molecular content, fusion transcripts will be handled like any other molecule in any subsequent TKSM modules.

### Single-cell barcoding

This module adds single-cell barcode sequences to the molecules using data generated by the Transcribing module. If the expression data passed to the Transcribing module was generated in single-cell expression mode, the Transcribing module will add the cellular barcode information of each molecule in a special tag (e.g. `CB:ACTACGAAGAAACCAT`) in its MDF output. These cellular barcode tags are mainly used by the Single-cell Barcoding module to add the barcode sequence to their respective molecules.

### Tagging

The Tagging module inserts custom sequences to the simulated molecules. This enables the user to add any combination of primers and/or UMI tags to the simulated molecules. The Tagging module is flexible and accepts IUPAC formatted strings of nucleotide codes to be appended at either the 5′ or 3′ ends of the molecules. For example, `tksm tagging -3 AYNN` will append at the start of each molecule a random tag that begins with `A`, followed by `C` or `T`, followed by 2 random nucleotides.

### Filtering

The Filtering module takes a series of conditions on the molecule records and filters any molecules that fail one or more of these conditions. The module supports conditions on the length of the molecule, overlaps with genomic loci or chromosomes, and the presence of specific tags (e.g. cellular barcode). This module is useful for creating different pipelines for the molecules that pass the filter and those that fail it.

### PCR

The PCR module duplicates the input material simulating the PCR process in a manner similar to work done in Calib [Orabi et al., 2019] and Minnow [Sarkar et al., 2019]. The PCR module takes as parameters the number of PCR cycles, *c*, the error rate per duplicated base, *e*, and the PCR efficiency, *f ∈* [0, 1]. Additionally, the PCR module takes the desired number of molecules, *N*, to be selected from the exponentially many molecules that will be present at the end of the PCR process. Conceptually, in each PCR cycle, the module randomly selects a set of molecules to duplicate equal to the number of input molecules multiplied by *f*. It then inserts random substitution errors to the duplicated molecules equal to their total length multiplied by *e*. It then proceeds to the next cycle using the old and new sets of molecules as input. Finally, from the exponentially many molecules created, the modules randomly outputs *N* of them. The parameter *N* is generally a small fraction of the the number of molecules present at the end of the PCR process, which is equal to *M ×* (1 + *f*)^*c*^ where *M* is the number of input molecules. Therefore, creating all the PCR molecules at the same time would lead to a huge memory footprint. Rather, TKSM processes each molecule independently by creating a truncated duplication tree for it. Initially, the tree has a single node representing the original molecule. In each PCR cycle, the module decides randomly (with success rate of *f*) for each node in the tree if this node should be expanded into a subtree. If the expansion test is successful, TKSM adds a new child to the node, associated to a newly duplicated molecule obtained from its parent by simulating a number of random mutations equal to the molecule length multiplied by *e*. Then TKSM decides whether to capture (i.e. output) this newly created molecule randomly with success rate equal to probability of capturing a molecule in the output: *N/ M ×* (1 + *f*)^*c*^.

### polyA tails addition

The polyA module appends a polyA tail to the molecules in its input MDF following a normal distribution of the length of the tails with a specified mean and standard deviation. To estimate these parameters from a real dataset, we use a simple script described by Orabi et al. [2023] to detect the length of the polyA tails of the reads of the real dataset.

### Strand flipping

This module takes an MDF and randomly reverses the order of the intervals of the molecules in the MDF according to a user-defined probability *p*. If a molecule is reversed, the module then adds a tag to its intervals indicating that their sequence is reversed complemented. The module is useful for simulating sequencing protocols that are not strand-specific.

### Gluing

The Gluing modules takes an MDF and a user specified probability *p*. It then processes each molecule, and with probability 1 − *p* it outputs the molecule with no modification and with probability *p* it prepends the molecule to the next molecule. The Gluing module aims to simulates the behaviour of ONT signal mis-segmentation process in which sometimes the ONT software fails to segment the signal of two consecutive molecules and as a result outputs their sequences as a single read.

### Shuffling

This module takes an MDF and outputs its molecules in random order. To allow for reduced output latency and memory consumption, the module buffers the input into a dynamically allocated array with a maximum size of *N* (user defined with default of *N* = ∞). Once the buffer array is full, the module will randomly output one of its molecules and replace it with a new incoming molecule. Once the input is exhausted, the buffer array is shuffled in-place and its molecules are outputted. The smaller *N* is, the more localized the shuffling will be. The randomized shuffling of the molecules enables the user to generate a random subsample from an MDF and is necessary for modules that assume a random order of the molecules such as the Gluing module.

### Truncating

This module simulates the process by which only a portion of the molecule is sequenced due to truncation. Such sequence truncation is caused by library preparation artifacts or, specifically in the case of ONT sequencing, by early stopping of the sequencing due to pore blocking [Soneson et al., 2019, Amarasinghe et al., 2020b]. To simulate the truncation of transcriptomic molecules, we use a two-dimensional kernel density estimation (KDE) model to decide the truncation length with respect to the transcript length, similar to Trans-Nanosim [Hafezqorani et al., 2020]. In TKSM, we use Scikit-learn’s KDE implementation [Pedregosa et al., 2011]. The exact details of how the KDE models are derived and the differences between our approach and that of Trans-Nanosim are described in Section S1.5 of the Supplementary Material.

### Merging

The Merging module is a simple module that concatenates the MDF outputs of one or more other TKSM modules into a single MDF file. This module is another entry-point module alongside the Transcribing module. The main use of this module is to enable simulation pipelines that take as input the output of one or more simulation pipelines; for example the user may use this module to build a mixed-sample simulation dataset by merging the MDF output of multiple pipelines that use the Transcribing modules on different samples.

In the piped version of TKSM, the Merging module is slightly more complicated than a simple simple concatenation operation. This is because concatenating files linearly and in-order can result in a deadlock when the input and output of the modules is piped. To avoid such deadlock, we implemented a multi-threaded version of the Merging module that assigned a CPU thread to each file, concurrently consuming the input files.

### Sequencing

The Sequencing module takes an MDF and the genome reference file(s) of the simulated sample and generates sequencing reads of the molecules in the MDF. The module can generate perfect reads (i.e. with no base-level errors) or erroneous reads, under a given error model. To generate erroneous reads we integrated some functions from the Badread [Wick, 2019] long-read simulator into a multi-threaded implementation in the Sequencing module. We also used Badread’s method of building a *k*-mer substitution error and quality score model. As part of TKSM Snakemake, the user may specify to train Badread models on the given real samples or to use pre-trained Badread models. This allows TKSM to be easily applied to new or future long-read sequencing chemistries.

## 2.2 Customizable TKSM pipelines using Snakemake

An important design choice for TKSM is to make it easily customizable by the user, i.e. to make it easy to build a, possibly complex, simulation pipeline using the modules described above. To achieve that, we packaged TKSM with Snakemake and configuration scripts that can be edited by the user to add new modules or to defined simulation experiments using any arrangement of TKSM modules. To define a simulation pipeline, the user lists the names of required TKSM modules and specify, for the modules that require model construction, the real samples to build such models on. Additionally, using the Merging module, the user may build complex pipelines that are composed of different linear pipelines. An example of the configuration script is presented in Supplementary Material Section S1.

## 3 Results

To illustrate TKSM and assess its performances, we designed three simulation pipelines to emulate examples of standard transcriptomic sequencing protocols. Specifically, we present simulations of a standard bulk RNA sequencing experiment, a hybrid long-short read single-cell RNA sequencing (scRNA-seq) experiment, and an RNA sequencing experiment similar to the bulk RNA sequencing experiment but with 100 random gene fusion events added. The Snakemake configuration files that specify these simulation pipelines are presented in Listings S2, S3, and S4 of the Supplementary Material. We use Matplotlib Hunter [2007] and Numpy [Harris et al., 2020] to generate many of our results.

In the standard bulk RNA-seq experiment, we primarily compare against Trans-Nanosim [Hafezqorani et al., 2020] and try to conform to its pipeline design using TKSM modules. For both the bulk and gene fusion experiments, we use an RNA-seq sample generated from the MCF7 cell line by Chen et al. [2021] (direct RNA, replicate 1, run 2). We first accessed the SG-NEx data on 2020-06-17 via https://registry.opendata.aws/sgnex/. For the scRNA-seq experiment, we used an in-house dataset, named N1, first described by Ebrahimi et al. [2022]. N1 follows the short-long single-cell hybrid protocol described previously in the literature [Singh et al., 2019, Tian et al., 2021, Gupta et al., 2018]. In this manuscript, we use a random subsample of N1 with ∼1M long-reads. The three TKSM pipelines are illustrated in Figure 2 and Figure S8 of the Supplementary Materials.

**Figure 2.**
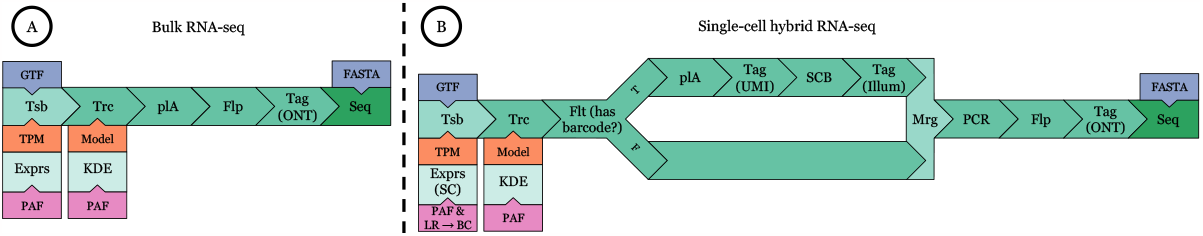
TKSM simulation pipelines. A) Typical RNA-seq simulation pipeline that imitates Trans-Nanosim’s workflow. B) Single-cell long-read simulation pipeline. The pipeline makes use of the Filtering and Merging modules to add the short-read Illumina adapter and 10x Genomics cellular barcodes only to molecules that have a tag indicating that they should have a cellular barcode.

Using these experiments, our goal is to assess TKSM on multiple metrics: i) the similarity of the simulated data to input real data on measures such as transcript expression, molecule sequence truncation and single cell barcode detection rates, and ii) time and memory footprint of various steps, iii) the ability to generate gene fusion events that can be detected by standard gene fusion tools. The results for the gene fusion experiment are presented in Section S1.7 of the Supplementary Material. Note that all the results presented in this section are reproducible using Snakemake scripts provided on the TKSM’s GitHub repository.

## 3.1 Data characteristics

Our goal in this section is to demonstrate that TKSM is capable of producing simulated sequencing data that has realistic characteristic in terms of its biological features (e.g. isoform expression) and technological artifacts (e.g. sequencing error). To assess these characteristics, we compare the simulated datasets to the input real dataset they were based on (MCF7 for bulk RNA-seq and N1 for scRNA-seq).

### Transcript expression profiling

We used LIQA [Hu et al., 2021] to compute the expression profiles of the different simulated and real bulk RNA-seq datasets. We additionally generated transcript expression profiles using Minimap2, trans-Nanosim, and TKSM. For Minimap2, we used the number of primary alignment of the long-reads to the transcripts as transcript counts. For trans-Nanosim and TKSM, we use the `read_analysis.py quantify` and `tksm abundance` commands, respectively, to generate transcript counts. We then normalize all the generated counts to be transcript-per-million (TPM) counts by dividing each transcript count by the total expression and multiplying by one million. For the scRNA-seq dataset, we used TKSM single-cell mode of the abundance estimation utility. We then combine the TPM counts of each gene since the number of unique single-cell barcode and transcript pairs is very large compared to the the number of generated reads. The TPM counts using these different methods on the bulk RNA-seq datasets are plotted in Figures 3 and S1 and for the scRNA-seq dataset in Figure 4.

**Figure 3.**
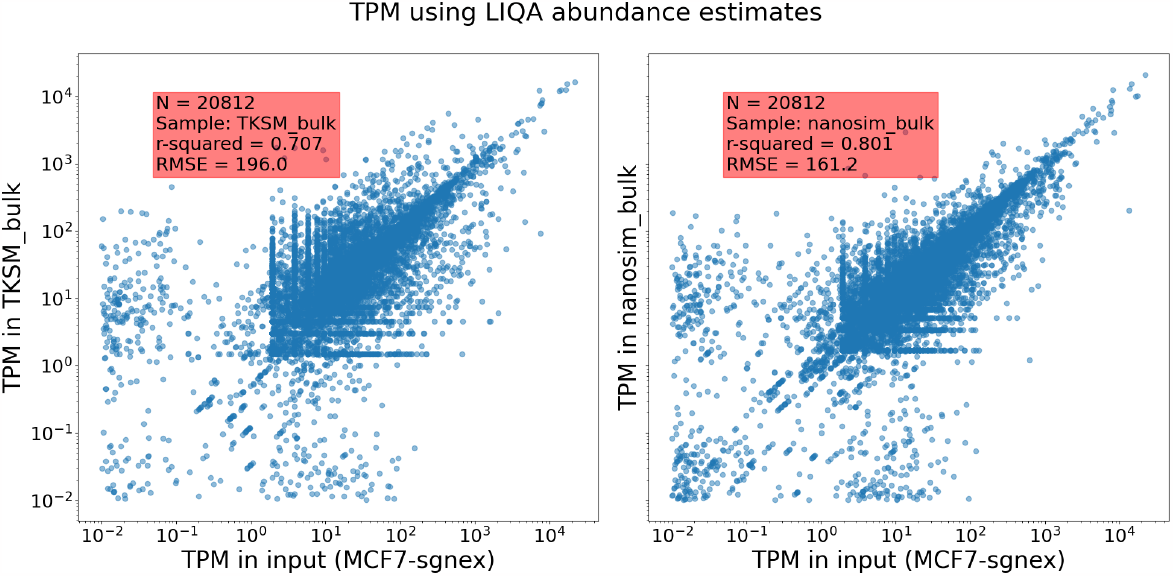
Transcript expression (TPM) of bulk RNA-seq datasets computed by LIQA.

**Figure 4.**
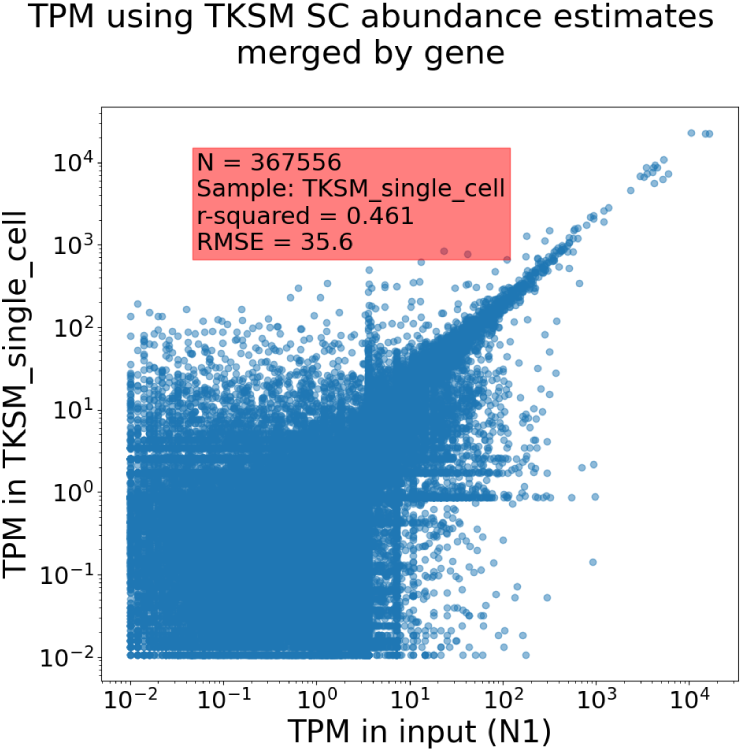
Hybrid scRNA-seq using TKSM single-cell transcript expression utility. Each point represents the expression sum of the transcripts of the same gene and cellular barcode.

On the bulk RNA-seq datasets, TKSM and Trans-Nanosim have high concordance with the input real dataset with correlation coefficients of 0.71 and 0.71, respectively. Both tools have small root mean square error (RMSE) rates of 196 and 161, respectively. On the scRNA-seq dataset, TKSM has a lower correlation coefficient of 0.46 compared to its performance on the RNA-seq dataset. However, a reduction in the correlation coefficient for the scRNA-seq dataset is expected since it includes over 367K gene/barcode data points compared and thus has a lot more opportunity for variation from the input counts.

### Truncation

To measure the truncation level of the sequencing pipeline, we mapped the long-reads of the different datasets to the transcriptome using Minimap2. For each primary alignment, we compute the read mapping length (*query_end* − *query_start*) which we use as a proxy for the post-truncation sequencing length. The mapping lengths are plotted as bar graphs in Figure 5 and Figure S2 of the Supplementary Material. Both Trans-Nanosim and TKSM generate very similar read length distribution compared to the input datasets.

**Figure 5.**
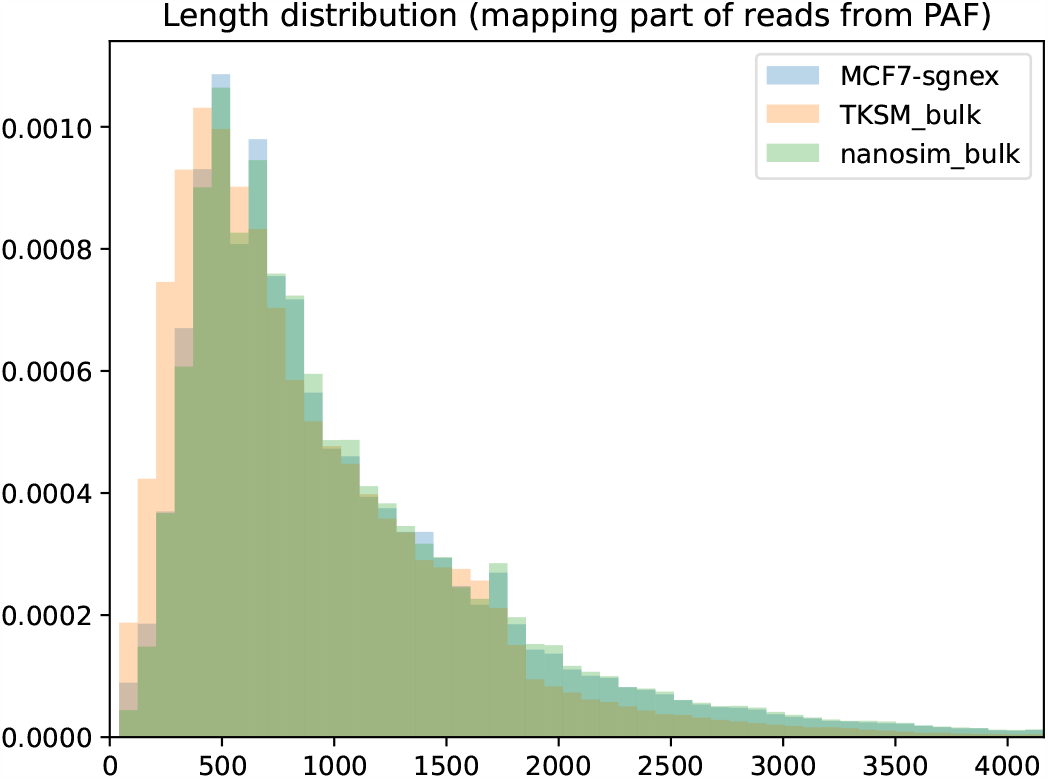
Mapped length of the reads in the bulk RNA-seq datasets vs the length of the transcript of their primary alignment using Minimap2.

### PolyA tail lengths

To compute the length of the observed polyA tails on the long-reads, we use a simple poly-A detection method described by Orabi et al. [2023]. The distribution of the polyA tails and lengths is plotted in Figure 6 for the bulk RNA-seq dataset and in Figure S3 of the Supplementary Material. The real N1 scRNA-seq dataset seem to have a bi-modal distribution (polyA tails *<* 10bp and ∼ 25bp) which are not both captured by the TKSM simulated dataset. However, the distribution observed for the TKSM dataset is closer to the real data distribution compared to Trans-Nanosim.

**Figure 6.**
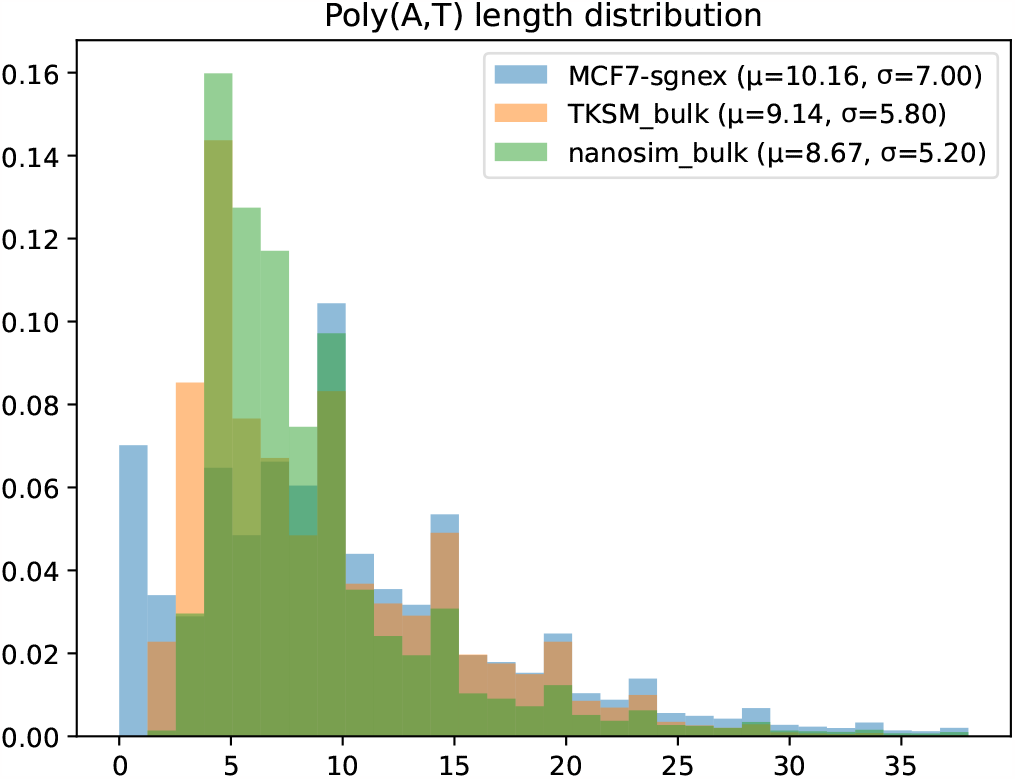
Poly-A length of the reads in the bulk RNA-seq datasets as computed by TKSM. TO SUPP. MAT.

### Base-level sequencing errors

To assess the base-level sequencing error profile of the datasets, we aligned the reads to the transcriptome using Minimap2 with `-c` flag to generate the alignment CIGAR strings. We then compute, as a percent of the mapped read length, the match, substitution, insertion, and deletion rates. The distributions of the sequencing errors are plotted in Figure S4 of the Supplementary Materials both the bulk RNA-seq and scRNA-seq datasets.

### Single-cell barcode profile

To assess the distribution of edit error on the cellular barcodes present on the long-reads, we performed pair-wise alignment of each whitelist cellular barcode to each long-read using Edlib [Šošić and Šikić, 2017] Python package. Instead of aligning to the whole long-read, we only considered the ranges [25, 75) and [−75, −25) of the long-reads to reduce the running time. The cellular barcodes whitelist was generated by running scTagger [Ebrahimi et al., 2022] on the full long-read dataset. We considered each cellular barcode and its reverse complement as independent barcodes. Only the alignments minimizing the edit distance, *d*, are kept from all the computed pair-wise alignments. If only one such alignment exists, then we consider it a unique alignment. Otherwise we consider the alignments to be ambiguous. The TKSM generated dataset has a realistic distribution of cellular barcode matching in terms of the ambiguity of assigning each long-read to a barcode and the number of errors detected on the barcode as shown in Figure 7.

**Figure 7.**
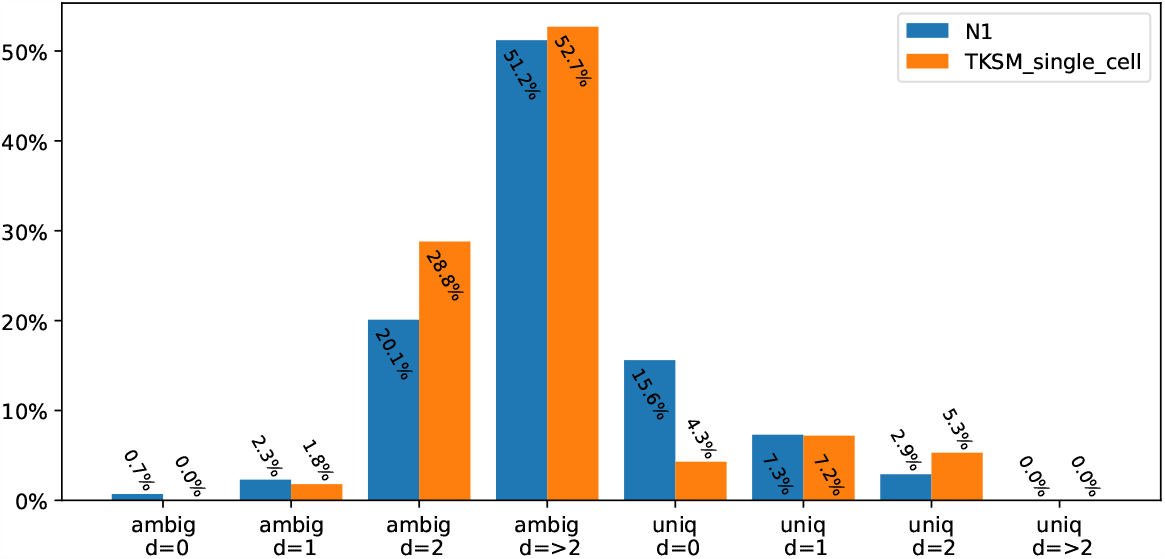
Brute-force detection of the cellular barcodes on a random sample of 50,000 long-reads from each dataset.

We also wanted to assess the distribution of where the Illumina short-read adapter is located on the the long-reads. To compute these loci, we ran scTagger on the scRNA-seq long-read datasets. We observe that the simulated dataset generated by TKSM has a similar distribution to the real dataset in terms of the loci of the SR adapter on the LRs and in terms of the number of LRs that scTagger is unable to detect a SR adapter on. This is demonstrated in Figure S5 of the Supplementary Materials.

## 3.2 Time and memory use

We used GNU Time to monitor the memory and CPU use of the tested tools. The overall results for the bulk RNA-seq and the scRNA-seq experiments are presented in Table 1. The detailed results for the different steps of all these experiments are presented in Tables S1 and S2 of the Supplementary Materials. For both tools, we separate the preprocessing steps (e.g. mapping, sequence error profiling, cell barcode detection) from the core steps (e.g. molecule generation, sequencing). For TKSM, we run its core processes twice: once in regular (i.e. blocking) mode and once in piped mode.

**Table 1.** Time and memory usage as reported by GNU Time (v1.9) for bulk RNA-seq and scRNA-seq datasets. All tools were allowed to run with up to 32 CPUs. For regular (i.e. blocking) TKSM runs, reported real and user times are the sum of individual processes and memory is the maximum memory use by any individual process. For piped TKSM runs, the reported time and memory values are captured by GNU Time for the whole pipeline treated as a single process.

| Experiment | Tool |  | Time (min) |  | Memory (GB) |
| --- | --- | --- | --- | --- | --- |
|  |  |  | Real | User |  |
| Bulk RNA | Nanosim | preprocess | 46.2 | 603.6 | 36.2 |
|  |  | core | 8.7 | 224.0 | 0.8 |
|  | TKSM | preprocess | 28.9 | 461.5 | 9.4 |
|  |  | core | 5.5 | 86.0 | 3.5 |
|  |  | pipelined core | 4.0 | 90.1 | 3.5 |
| scRNA | TKSM | preprocess | 17.8 | 227.3 | 12.5 |
|  |  | core | 16.2 | 283.2 | 3.5 |
|  |  | pipelined core | 12.2 | 340.4 | 3.5 |

As we observe in Table 1, TKSM finishes its preprocessing and core processing in, respectively, %36 and %37 less time than Trans-Nanosim despite generating the same amount of data and running similar pipelines. For the core processes, TKSM runs in 27% or 25% less time, respectively, on bulk RNA-seq and scRNA-seq datasets, when it runs in piped mode compared to its regular mode and without any increase in memory usage.

## 4 Conclusion

TKSM is a modular and high-performing transcriptomic LR sequencing simulator. Its modular design enables the user to construct a large verity of sequencing experiments with minimal effort. TKSM’s standardized input and output for its modules allows us and the users of TKSM to add new modules that target existing and future library preparation techniques that TKSM currently does not target. For example, it is easy to envision an alternative entry-point module to the Transcribing module that generates nucleic acid molecules from DNA fragmentation while still making use of the rest of TKSM modules. TKSM also performs well in terms of generating realistic datasets with characteristics matching the real datasets it is imitating. Additionally, TKSM is engineered with efficient CPU and memory use in mind and its performance on those metrics is excellent.

## Author contributions

Conceptualization: F.K. and B.O.; Methodology: F.K. and B.O.; Software: F.K. and B.O.; Formal analysis: F.K. and B.O.; Investigation: F.K. and B.O.; Resources: R.F and F.H.; Data Curation: F.K., B.O., R.F, and F.H.; Writing - Original Draft: F.K. and B.O.; Writing - Review & Editing: F.K., B.O., C.C., and F.H.; Visualization: F.K. and B.O.; Supervision: C.C. and F.H.; Project administration: C.C. and F.H.; Funding acquisition: B.O., C.C., and F.H.

## Data availability

TKSM is available as an open source software at https://github.com/vpc-ccg/tksm. Datasets used in this manuscript are publicly available via https://registry.opendata.aws/sgnex/ for the MCF7 dataset and at https://doi.org/10.6084/m9.figshare.23155145 for the N1 sample.

