## Supplementary Materials for "TKSM: Highly modular, user-customizable, and scalable transcriptomic sequencing long-read simulator"

### S1 Supplementary Material

#### S1.1 Codes listings

```
TS_experiments:
    head_1:
        pipeline:
            - Tsb:
                params: "--molecule-count 200000"
                model: "MCF7-sgnex"
                mode: Xpr
            - Trc:
                params: ""
                model: "MCF7-sgnex"
            - plA: {}
    head_2:
        pipeline:
            - Tsb:
                params: "--molecule-count 100000"
                model: "N1"
                mode: Xpr_sc
            - Trc:
                params: ""
                model: "N1"
            - plA: {}
            - SCB: {}
            - Tag:
                params: "--format5 10"
    experiment_1:
        pipeline:
            - Mrg:
                sources: ["head_1", "head_2"]
            - PCR:
                params: "--cycles 10 -x T4 --molecule-count 1000000"
            - Flp: {}
            - Seq:
                params: "--skip-qual-compute"
                model: "nanopore2020"
samples:
    "MCF7-sgnex":
        fastq:
            - data/samples/MCF7-sgnex.fastq.gz
        ref: Homo_sapiens
    "N1":
        fastq:
            - data/samples/N1.fastq
        ref: Homo_sapiens
refs:
    Homo_sapiens:
        cDNA: data/refs/Homo_sapiens.cdna.fa
        DNA: data/refs/Homo_sapiens.dna.fa
        GTF: data/refs/Homo_sapiens.gtf
barcodes:
    10x_bc: data/refs/3M-february-2018.txt.gz
```

**Listing S1.** An example for a TKSM Snakemake configuration file with a simulation pipeline that uses as input the outputs of two other simulation pipelines.

```
TS_experiments:
    bulk_RNA:
        pipeline:
            - Tsb:
                params: "--molecule-count 1000000"
                model: "MCF7-sgnex"
                mode: Xpr
            - Trc:
                params: ""
                model: "MCF7-sgnex"
            - plA:
                params: "--normal=15,7.5"
            - Flp:
                params: "-p 0.5"
            - Tag:
                params: "--format5 AATGTACTTCGTTCAGTTACGTATTGCT --format3 GCAATACGTAACTGAACGAAGT"
            - Seq:
                params: "--skip-qual-compute"
                model: "MCF7-sgnex"
```

**Listing S2.** TKSM Snakemake configuration file for the bulk RNA-seq simulation experiment.

```
TS_experiments:
    single_cell_head:
        pipeline:
            - Tsb:
                params: "--molecule-count 1000000"
                model: "N1"
                mode: Xpr_sc
            - Trc:
                params: ""
                model: "N1"
    single_cell_p1:
        pipeline:
            - Mrg:
                sources: ["single_cell_head",]
            - Flt:
                params: "-c \"info CB\""
            - plA:
                params: "--normal=15,7.5"
            - Tag:
                params: "--format3 10"
            - SCB:
                params: ""
            - Tag:
                params: "--format3 AGATCGGAAGAGCGTCGTGTAG"
    single_cell_p2:
        pipeline:
            - Mrg:
                sources: ["single_cell_head",]
            - Flt:
                params: "-c \"info CB\" --negate"
    single_cell:
        pipeline:
            - Mrg:
                sources: ["single_cell_p1", "single_cell_p2"]
            - PCR:
                params: "--cycles 5 --molecule-count 5000000"
            - Flp:
                params: "-p 0.5"
            - Tag:
                params: "--format5 AATGTACTTCGTTCAGTTACGTATTGCT --format3 GCAATACGTAACTGAACGAAGT"
            - Seq:
                params: "--skip-qual-compute"
                model: "N1"
```

**Listing S3.** TKSM Snakemake configuration file for the single-cell RNA-seq simulation experiment.

```
TS_experiments:
    gene_fusion_RNA:
        pipeline:
            - Tsb:
                params: "--molecule-count 1000000 --fusion-count 100"
                model: "MCF7-sgnex"
                mode: Xpr
            - Trc:
                params: ""
                model: "MCF7-sgnex"
            - plA:
                params: "--normal=15,7.5"
            - PCR:
                params: "--cycles 5 --molecule-count 5000000"
            - Flp:
                params: "-p 0.5"
            - Tag:
                params: "--format5 AATGTACTTCGTTCAGTTACGTATTGCT --format3 GCAATACGTAACTGAACGAAGT"
            - Seq:
                params: "--skip-qual-compute"
                model: "MCF7-sgnex"
```

**Listing S4.** TKSM Snakemake configuration file for the bulk RNA-seq simulation experiment with 100 random gene fusion events added (option `--fusion-count 100` of the `Tsb` module).

#### S1.2 Figures

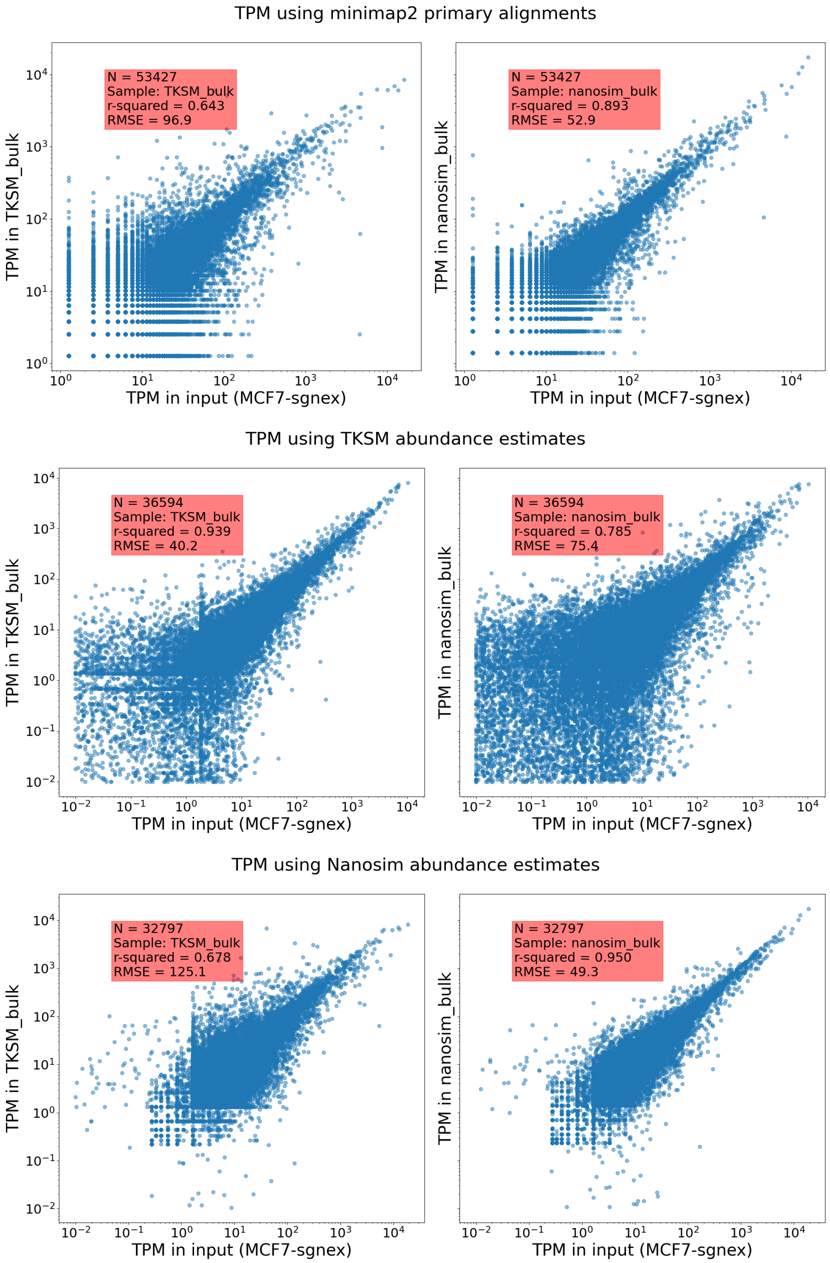


**Figure S1.** Bulk RNA-seq transcript/gene expression using different methods.

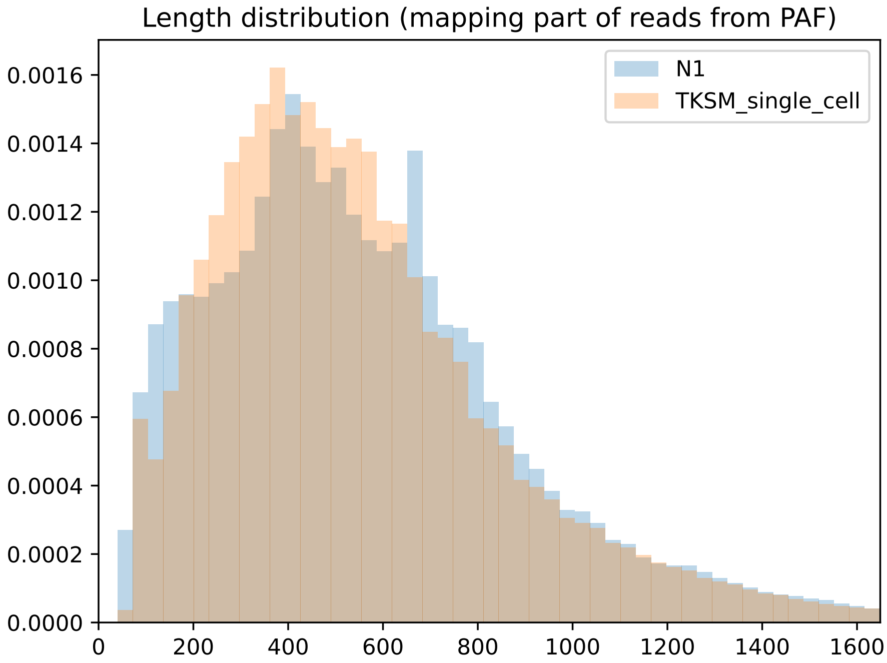


**Figure S2.** Mapped length of the reads in the scRNA-seq datasets vs the length of the transcript of their primary alignment using Minimap2.

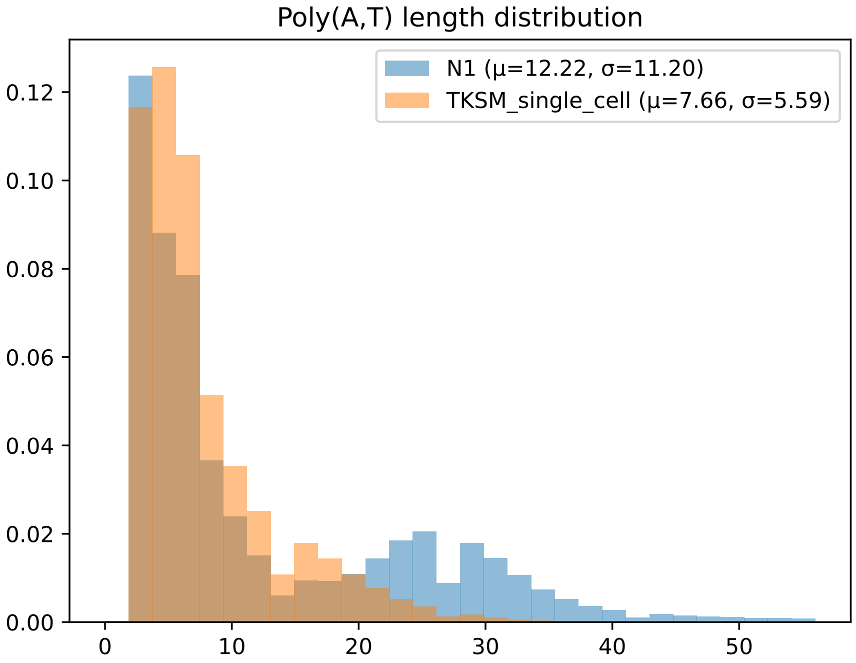


**Figure S3.** Poly-A length of the reads in the scRNA-seq datasets as computed by TKSM.

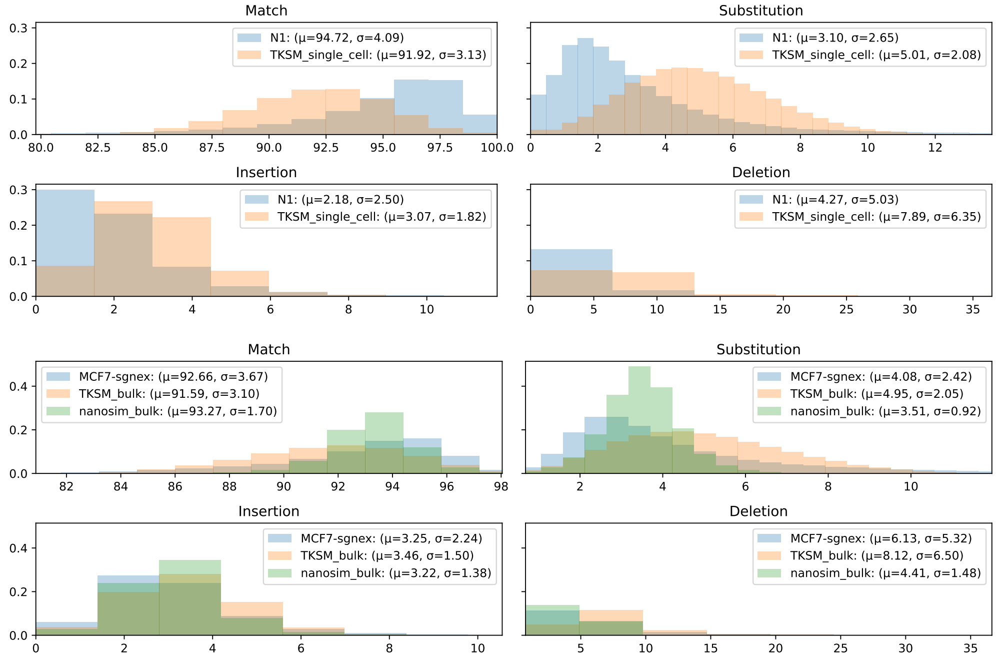


**Figure S4.** Substitution, insertion, and deletion base-level error rates using the CIGAR strings of Minimap2 mapped reads of the bulk RNA-seq (top two rows) and scRNA-seq datasets (bottom two rows).

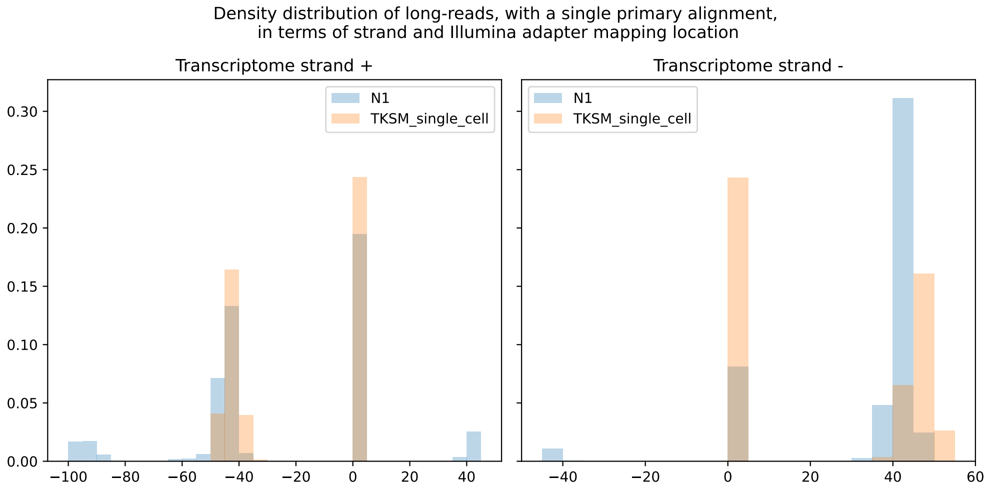


**Figure S5.** Distribution of the detected Illumina short-read adapter on the long-reads separated by the transcript strand to which the long-read maps to. The $x$-axis represent the match location of the Illumina adapter on the long-read. Negative $x$-axis values represent match location from the end of the read for Illumina adapters that map on the negative strand of the long-read. The 0 point on the $x$-axis represent long-reads that have no Illumina adapter detected on them.

#### S1.3 Tables

**Table S1.** Bulk RNA-seq

| Tool | Process | Real time (min) | User time (min) | Memory (GB) | CPUs |
|---|---|---|---|---|---|
| Nanosim (preprocess) | `NS_analysis` | 26.4 | 538.1 | 36.2 | 32 |
| | `NS_quantify` | 19.8 | 65.4 | 6.8 | 32 |
| | **Total** | **46.2** | **603.6** | **36.2** | **32** |
| Nanosim (simulate) | **Total** | **8.7** | **224.0** | **0.8** | **32** |
| TKSM (preprocess) | `minimap2` | 6.7 | 169.1 | 9.4 | 32 |
| | `abundance` | 1.5 | 1.4 | 3.8 | 1 |
| | `model_truncation` | 0.4 | 3.0 | 0.2 | 32 |
| | `minimap2 (badread)` | 2.2 | 54.9 | 5.8 | 32 |
| | `badread_error_model` | 1.8 | 1.7 | 4.9 | 1 |
| | `badread_qscore_model` | 7.5 | 7.4 | 4.9 | 1 |
| | **Total** | **28.9** | **461.5** | **9.4** | **32** |
| TKSM (core) | `transcribe` | 1.3 | 1.2 | 3.5 | 1 |
| | `truncate` | 0.2 | 0.2 | 0.0 | 1 |
| | `polyA` | 0.3 | 0.3 | 0.0 | 1 |
| | `flip` | 0.3 | 0.3 | 0.0 | 1 |
| | `tag` | 0.4 | 0.4 | 0.0 | 1 |
| | `sequence` | 3.1 | 83.7 | 3.3 | 32 |
| | **Total** | **5.5** | **86.0** | **3.5** | **32** |
| TKSM (core;piped) | `transcribe` | 3.9 | 1.1 | 3.5 | 1 |
| | `truncate` | 3.9 | 0.4 | 0.0 | 1 |
| | `polyA` | 3.9 | 0.5 | 0.0 | 1 |
| | `flip` | 3.9 | 0.5 | 0.0 | 1 |
| | `tag` | 3.9 | 0.6 | 0.0 | 1 |
| | `sequence` | 3.9 | 87.1 | 3.2 | 32 |
| | **Total** | **4.0** | **90.1** | **3.5** | **32** |

**Table S2.** Hybrid scRNA-seq

| Tool | Pipe path | Process | Real time (min) | User time (min) | Memory (GB) | CPUs |
|---|---|---|---|---|---|---|
| TKSM (preprocess) | | scTagger (extract BC) | 1.0 | 0.9 | 3.0 | 1 |
| | | scTagger (LR seg) | 0.8 | 2.5 | 1.5 | 32 |
| | | scTagger (match) | 6.7 | 159.7 | 0.9 | 32 |
| | | minimap2 (badread) | 1.0 | 0.9 | 3.2 | 32 |
| | | badread_error_model | 1.3 | 1.1 | 12.5 | 1 |
| | | badread_qual_model | 3.4 | 3.3 | 3.1 | 1 |
| | | minimap2 | 2.4 | 55.8 | 6.2 | 32 |
| | | model_truncation | 0.2 | 2.3 | 0.2 | 32 |
| | | abundance (SC) | 1.1 | 1.0 | 2.8 | 1 |
| | | **Total** | **17.8** | **227.3** | **12.5** | **32** |
| TKSM (core) | head | transcribe | 1.2 | 1.2 | 3.5 | 1 |
| | | truncate | 0.3 | 0.3 | 0.0 | 1 |
| | path_1 | filter | 0.2 | 0.2 | 0.0 | 1 |
| | | polyA | 0.1 | 0.1 | 0.0 | 1 |
| | | tag | 0.2 | 0.2 | 0.0 | 1 |
| | | single-cell barcode | 0.2 | 0.2 | 0.0 | 1 |
| | | tag | 0.2 | 0.2 | 0.0 | 1 |
| | path_2 | filter | 0.2 | 0.2 | 0.0 | 1 |
| | tail | PCR | 1.3 | 1.2 | 1.4 | 1 |
| | | flip | 1.4 | 1.3 | 0.0 | 1 |
| | | tag | 1.8 | 1.7 | 0.0 | 1 |
| | | sequence | 9.2 | 276.5 | 3.3 | 32 |
| | | **Total** | **16.2** | **283.2** | **3.5** | **32** |
| TKSM (core; piped) | head | transcribe | 1.7 | 1.4 | 3.5 | 1 |
| | | truncate | 1.7 | 0.3 | 0.0 | 1 |
| | path_1 | filter | 1.7 | 0.2 | 0.0 | 1 |
| | | polyA | 1.7 | 0.2 | 0.0 | 1 |
| | | tag | 1.7 | 0.2 | 0.0 | 1 |
| | | single-cell barcode | 1.7 | 0.2 | 0.0 | 1 |
| | | tag | 1.7 | 0.2 | 0.0 | 1 |
| | path_2 | filter | 1.7 | 0.2 | 0.0 | 1 |
| | tail | PCR | 12.2 | 2.2 | 1.3 | 1 |
| | | flip | 12.2 | 2.2 | 0.0 | 1 |
| | | tag | 12.2 | 2.8 | 0.0 | 1 |
| | | sequence | 12.2 | 325.2 | 3.3 | 32 |
| | | **Total** | **12.2** | **340.4** | **3.5** | **32** |

#### S1.4 Molecule Description Format

Each MDF entry (A molecule description) begins with a header line which consists of `+` symbol, followed by `<molecule_id>`, `<molecule_count>` and `<molecule_info>`:

```
+molecule_1 1 info1=1,2,3,4;info2;
```

This molecule header line is parsed by TKSM to:

```
{
 "Id": "molecule_1",
 "depth": 1,
 "info": {
 "info1": [1, 2, 3, 4],
 "info2": "."
    }
}
```

The header line is followed by a variable number of interval lines which are quite similar to the BED format:

```
chr start end orientation mods
```

The fields are tab-separated:

- The `chr` field is the name of the contig. Different intervals of the same molecule can be on different contigs.
- The `start` and `end` fields are the start and end positions of the interval (0-based, end-exclusive).
- The `orientation` field is the orientation of the interval (`+` or `-`). Different intervals of the same molecule can have different orientations.
- The `<mods>` field can be empty but the tab character preceding it is required. The `<mods>` is a list of comma separated base substitutions (no indels) local to the interval sequence represented in the current line. The substitutions are applied to the interval sequence before the strand is flipped (if the strand is `-`).

For example, consider the following FASTA contig and MDF entry:

```
FASTA:
>chr1
AGTCCCGTAA

MDF:
+m1 1
chr1 0 4 + 2C,3T
chr1 6 9 + 1G
```

Given the FASTA and MDF records, we can construct the sequence of the `m1` molecule.

- First we construct the sequence of the first interval: `(chr1, 0, 4) = AGTC`.
- We apply the modifications on positions 2 and 3: `AGTC -> AGCC -> AGCT`.
- Then we construct the second interval: `(chr1, 6, 9) = GTA`.
- We apply the modification on position 1: `GTA -> GGA`.

- Finally, we concatenate the two intervals to get the sequence of the `m1` molecule: `AGCTGGA`.

If the contig name is not in the provided reference FASTA but is nonetheless a valid nucleic sequence, TKSM will use the contig name as the contig sequence. Consider for example the following FASTA and MDF records:

```
FASTA:
>chr1
AGTC

MDF:
+m1 1
TT 0 2 +
chr1 0 4 +
```

Given the FASTA and MDF records, we can construct the sequence of the `m1` molecule.

- First we construct the sequence of the first interval. Since `TT` is not a reference name in the FASTA file, we treat it as a sequence itself: `("TT", 0, 2) = TT`.
- We construct the sequence of the second interval: `(chr1, 0, 4) = AGTC`.
- Finally, we concatenate the two intervals to get the sequence of the `m1` molecule: `TTAGTC`.

#### S1.5 Truncation Model Method

Regardless of the source of the truncation, TKSM models the truncation process as a function $f(X) = Y$ where the truncated length of the molecule, $Y$, is a function of the full length of the molecule, $X$. When estimating $f$, TKSM uses the mapping of real transcriptomic reads to the set of reference transcripts. In particular, the length of the transcript is taken as $X$ while the alignment length of the read on the transcript it originates from is taken as $Y$. A similar truncation model is used in Trans-Nanosim [Hafezqorani et al., 2020].

The Truncating module uses a kernel density estimation (KDE) method to build the truncation function $f$. To achieve this, we feed the pairs of transcript length and read alignment length to a two-dimensional kernel and then fit the KDE model to the input data. In TKSM, we use the scikit-learn [Pedregosa et al., 2011] implementation of KDE. In scikit-learn implementation, we can sample the KDE model to get $(X, Y)$ pairs. However, for the purpose of the truncation module, we need to sample the KDE for a $Y$ (truncated length) value given an $X$ value (molecule length).

To enable sampling the KDE model for $Y$ given $X$, we compute the likelihood of a large predefined set of $(X, Y)$ pairs. Specifically, we use the `score_samples` function of scikit-learn on $50 \times 50$ $(X, Y)$ pairs in the range $0 : 100 : 5000 \times 0 : 100 : 5000$. Then, given an $X$ value we find its closest lattice $X$-point and extract that $X$-point's column of $Y$ likelihood scores. We then sample the $Y$ values weighted by their likelihood scores and use the sampled $Y$ as the value for the truncation function $f(X)$.

Now to sample from this set of CDF, the following algorithm is used:

**Algorithm S1** TKSM truncation KDE sampling method.

$Input : X, Y, CDFs, y$

$y' \leftarrow lower\_bound(Y, y)$

$uni\_rand \leftarrow uniform\_random(0, 1)$

$pos1 \leftarrow lower\_bound(CDFs[y'], uni\_rand)$

$pos2 \leftarrow lower\_bound(CDFs[y' + 1], uni\_rand)$

$return\ (X[pos1] * (y - y') + X[pos2] * (y' + 1 - y))/2$

Where $X$ and $Y$ are the ranges of $X$-axis and $Y$-axis on the 2D Cartesian grid that is used to compute likelihoods. $CDFs$ is an array of CDFs for each point of $Y$. $uniform\_random$ is a random real number generator. $lower\_bound(A, a)$ is a binary search function that returns the position of the first number in $A$ that is not less than $a$. Finally, by taking the weighted average of random numbers of the closest two positions on the grid, we smooth the generated values.

Since the KDE model building procedure is computationally intensive, we implemented it in TKSM preprocessing utility, which computes and saves the lattice likelihood values. The pre-built models can then be used by the Truncating module.

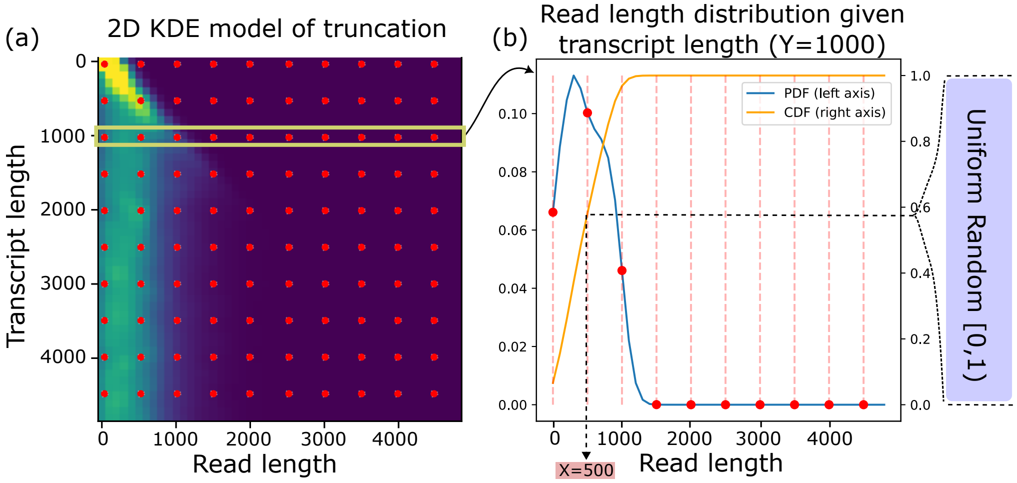


**Figure S6.** KDE Truncation method. a) Grid of values (500 bases apart in this example, but is more frequent in practice) are selected and likelihoods of each (x,y) pair on the KDE model are computed. To generate truncated read lengths for specific transcript lengths, TKSM takes a slice of likelihood values of the appropriate size. After normalization, these likelihoods are used as an estimation of a PMF and CDF is computed with a cumulative sum. b) Then TKSM converts uniform random values to random values on this distribution by finding lower bounds on CDF values using binary search (Weighted average strategy shown in the algorithm S1 is omitted for clarity) .

#### S1.6 Fusion simulation details

Fusion simulation is implemented as a sub-module to the transcribe module and activated when any fusion-related parameter is passed to the transcriber. This submodule can be summarized in three steps; characterization, configuration and modification.

The characterization step aims to define a list of genomic events that will induce gene fusion transcripts. Genomic events in this context are defined by two breakpoints (head and tail where fusion transcripts pass head before the tail) with contig, position and orientation and event rate which can be 1 for homozygous, 0.5 for heterozygous events and any value between 1 and 0 to indicate fusion population in somatic samples. These events are indexed for efficient access in the latter steps. The result of this step is a list of pairs of breakpoints, each pair being composed of one breakpoint within each of the tail and head gene of the fusion. While biologically it is possible to have overlapping events (resulting in complex fusions), for the sake of clarity TKSM only simulates non-overlapping events.

The configuration step takes the gene annotation (GTF) and transcript abundances passed to the transcriber and the genomic events generated in the prior step. For each genomic event, transcripts overlapping with the breakpoints are gathered. Fusion transcripts are determined by the set of transcript pairs generated by the Cartesian product of the head-overlapping and tail-overlapping transcripts (Figure S7). Total fusion expression is set by the sum of head transcript expression. This total expression is distributed across the fusion transcripts by following the Cartesian product of head-overlapping and tail-overlapping transcript expressions. Fusion transcript sequences are determined by concatenating the head exons before and tail exons after the breakpoints. Equivalent fusion transcripts are merged and their expression values are accumulated.

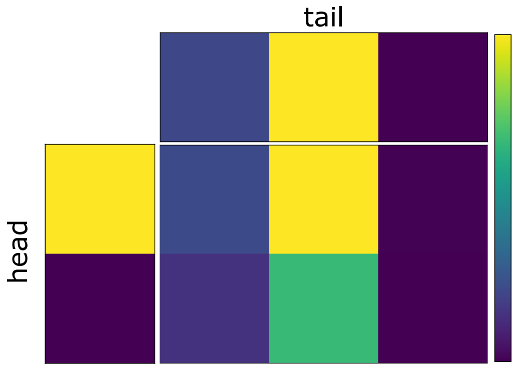


**Figure S7.** Cartesian product to decide fusion transcript expression. Each cell represent a transcript colored for its expression values (blue to yellow). In this example two genes with 2 and 3 transcripts are fused resulting 6 fusion transcripts.

Finally, the modification step adds the generated fusion transcript annotations and abundances to the simulation. Then, it modifies the expressions of the transcripts overlapping with the generated fusion events (on breakpoints or anything in between for deletions), reducing it with respect to the rate of the event.

**Algorithm S2** TKSM algorithm for generating random gene fusion events.

Input: Genes, FusionCount
$Genes \leftarrow shuffled(Genes)$
$G \leftarrow Genes[: 2 \times FusionCount]$ # *Take first 2 * FusionCount genes on shuffled Gene list*
$G \leftarrow sorted(g)$ #*by the genomic positions*
$fusion\_events \leftarrow list()$
**for** each consecutive ($g_1$,$g_2$) in G **do**
    $b_1 \leftarrow$ randomInt($g_1$.start, $g_1$.end)
    $b_2 \leftarrow$ randomInt($g_2$.start, $g_2$.end)
    fusion_events.add(FusionEvent($g_1$,$g_2$, $b_1$, $b_2$))
**end for**

**Algorithm S3** TKSM algorithm for generating fusion isoforms.

Transcripts, fusion_events
**for** each event in $fusion\_events$ **do**
    $t\_head \leftarrow Transcripts.overlaps(event.g_1)$
    $t\_tail \leftarrow Transcripts.overlaps(event.g_2)$
    $head\_count \leftarrow sum(t\_head.counts)$
    $tail\_count \leftarrow sum(t\_tail.counts)$
    $fusion\_isoforms \leftarrow list()$
    **for** each $t_1$ in $t\_head$ **do**
        **for** each $t_2$ in $t\_tail$ **do**
            $fusion\_count \leftarrow head\_count \times \frac{t_1.count}{head\_count} \times \frac{t_2.count}{tail\_count}$
            $iso_1 \leftarrow t_1.truncate(t_1.start, event.b_1)$
            $iso_2 \leftarrow t_2.truncate(t_2.start, event.b_2)$
            $fusion\_isoforms.add(concat(iso_1, iso_2), fusion\_count)$
        **end for**
    **end for**
**end for**

#### S1.7 Results of gene fusion experiment

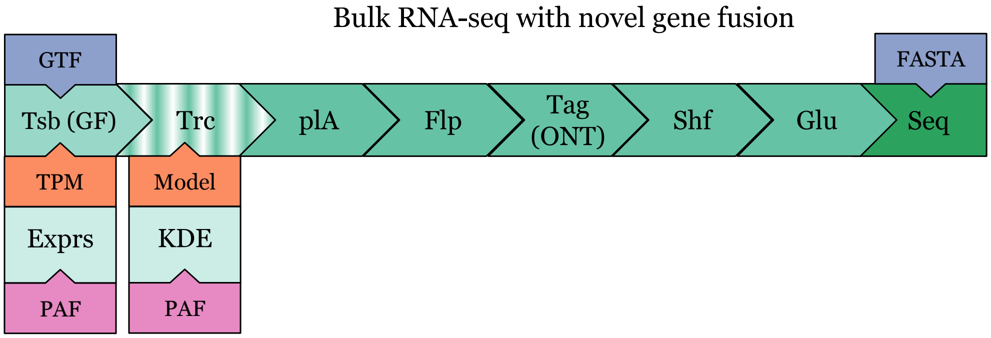


**Figure S8.** A TKSM RNA-seq pipeline that creates novel gene fusion events using the Transcribing module. The Glu module creates mis-segmentation errors resulting in chimeric reads. We also simulated datasets with and without truncation to evaluate the effect of truncation on the accuracy of gene fusion discovery.

Using TKSM, we simulated 50 random gene fusions, i.e., gene fusions generated from randomly selected gene pairs located on the same chromosome. We simulated datasets with and without truncation and with several mis-segmentation rates (0%, 1%, 2%, 4%, and 8%). Figure S8 illustrates the TKSM pipeline we designed for this gene fusion experiment. To test these datasets, we used gene fusion discovery tools Genion [Karaoglanoglu et al., 2022] and LongGF [Liu et al., 2020].

We observed that truncation negatively affect gene fusion detection. Note that our simulation pipeline is not strand-specific, i.e., molecules are sequenced from either end with equal probability. In this experiment, the inclusion of the truncation raised the number of undetectable fusions by either tool from 7 to 18 (Figure S9). While LongGF was resilient to the mis-segmentation, we observed a drop in the recall rate of the Genion especially when truncation was introduced to the dataset (Table S3). Additionally, when the mis-segmentation rate increases from 0% to 8%, the number of candidate gene fusions detected by Genion but subsequently discarded by one of its filters increase $\sim 5\times$. These findings show that TKSM is able generate gene fusion targets as well as un-segmentation errors that have direct effect on the running of gene fusion detection tools.

| Truncation | Glue rate | Tool | # | TP | FP | FN | F1 | Truncation | Glue rate | Tool | # | TP | FP | FN | F1 |
|---|---|---|---|---|---|---|---|---|---|---|---|---|---|---|---|
| FALSE | 0% | LongGF | 39 | 38 | 1 | 12 | 0.85 | TRUE | 0% | LongGF | 34 | 30 | 4 | 20 | 0.71 |
| | | Genion | 44 | 42 | 2 | 8 | 0.89 | | | Genion | 34 | 31 | 3 | 19 | 0.74 |
| | 1% | LongGF | 43 | 42 | 1 | 8 | 0.9 | | 1% | LongGF | 33 | 30 | 3 | 20 | 0.72 |
| | | Genion | 43 | 41 | 2 | 9 | 0.88 | | | Genion | 32 | 30 | 2 | 20 | 0.73 |
| | 2% | LongGF | 43 | 42 | 1 | 8 | 0.9 | | 2% | LongGF | 33 | 30 | 3 | 20 | 0.72 |
| | | Genion | 42 | 41 | 1 | 9 | 0.89 | | | Genion | 31 | 30 | 1 | 20 | 0.74 |
| | 4% | LongGF | 44 | 42 | 2 | 8 | 0.89 | | 4% | LongGF | 31 | 29 | 2 | 21 | 0.72 |
| | | Genion | 41 | 40 | 1 | 10 | 0.88 | | | Genion | 29 | 27 | 2 | 23 | 0.68 |
| | 8% | LongGF | 42 | 41 | 1 | 9 | 0.89 | | 8% | LongGF | 31 | 29 | 2 | 21 | 0.72 |
| | | Genion | 41 | 40 | 1 | 10 | 0.88 | | | Genion | 23 | 23 | 0 | 27 | 0.63 |

**Table S3.** Gene fusion detection results on the output of TKSM. For all eight experiments, the same 50 gene fusions were simulated.

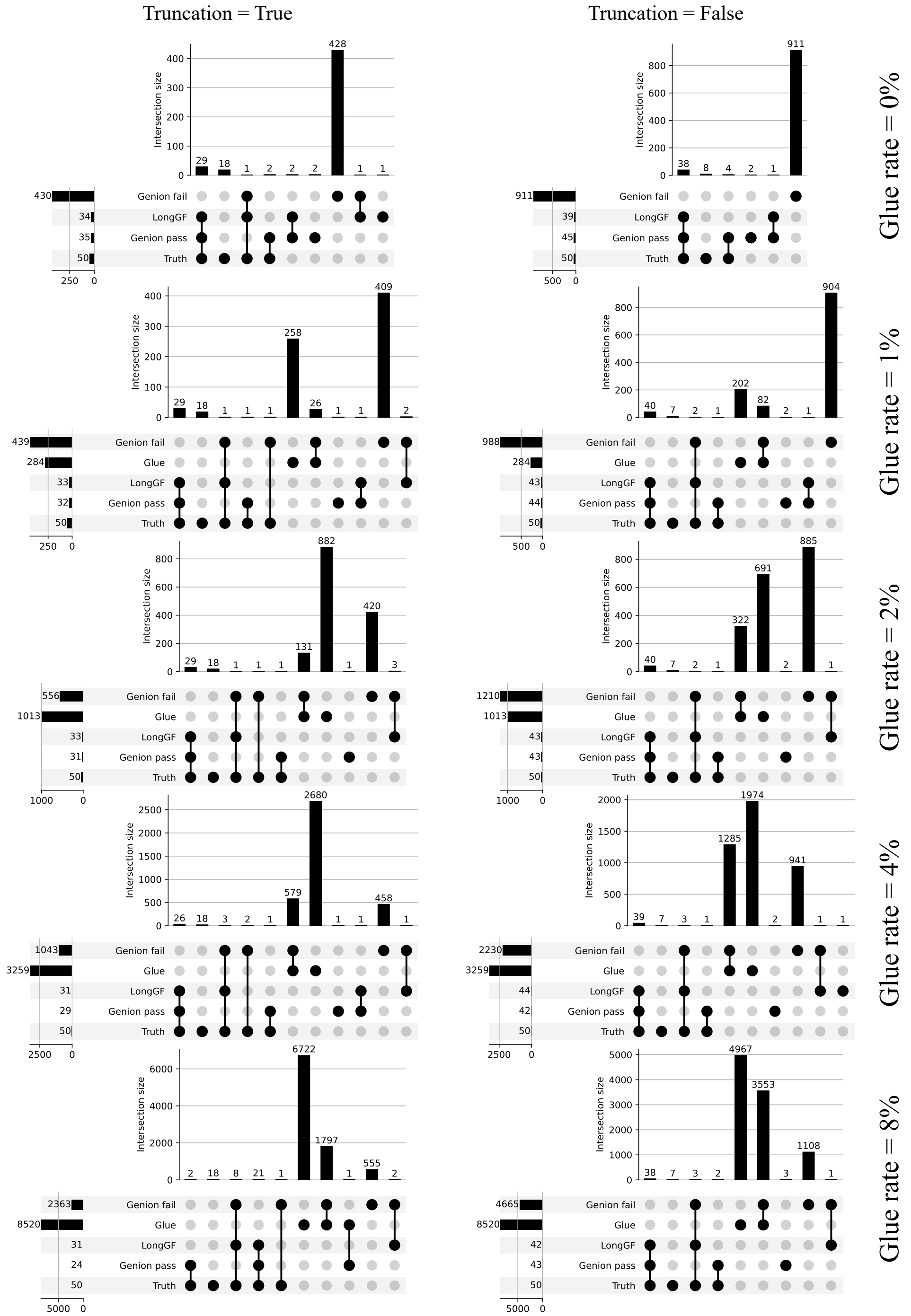


**Figure S9.** UpSet plots of the gene fusions simulated by TKSM or detected by Genion or LongGF. Note that UpSet plots are essentially Venn diagrams represented using bar graphs. For Genion, we report both the gene fusion detected by the tool (Genion pass) and those detected by Genion initially but later filtered by one of its filters (Genion fail). For the gene fusions simulated by TKSM, we separate between true gene fusions generated the Transcribing module (Truth) and false gene fusions that are generated by the Gluing module when it glues molecules of different genes (Glue). Note that for false gene fusions generated by the Gluing module, we require at least three molecules supporting each gene fusion to consider it in these plots.
